# Chimeric protein EWS-FLI1 drives cell proliferation in Ewing Sarcoma *via* overexpression of *KCNN1*

**DOI:** 10.1101/2023.04.24.538050

**Authors:** Maryne Dupuy, Maxime Gueguinou, Anaïs Postec, Régis Brion, Robel Tesfaye, Mathilde Mullard, Laura Regnier, Jérôme Amiaud, Marie Potier-Cartereau, Aurélie Chantôme, Bénédicte Brounais-Le Royer, Marc Baud’huin, Steven Georges, François Lamoureux, Benjamin Ory, Olivier Delattre, Françoise Rédini, Christophe Vandier, Franck Verrecchia

**Affiliations:** Nantes Université, Inserm UMR 1307, CNRS UMR 6075, Université d’Angers, CRCI2NA, F-44000 Nantes, France; N2C UMR 1069, University of Tours, INSERM, Tours, France; INSERM U830, Diversity and Plasticity of Childhood Tumors Lab, PSL Research University, SIREDO Oncology Center, Institut Curie, Paris, France

**Keywords:** Calcium / Ewing sarcoma / KCNN1 / Potassium channel / Proliferation

## Abstract

Ewing sarcoma (ES) is characterized by chimeric fusion proteins, which act as oncogenes. Over the last decade, patient survival has not increased, especially for high risk patients. Knowing that ion channels are studied for their implication in tumorigenesis, the aim of this work is to study the involvement of the SK1 potassium channels in ES. RNA-Seq analyses showed a high restricted expression of *KCNN1*, the gene encoding SK1, only in ES patients, and its expression is inversely correlated with patient survival. EWS-FLI1 silencing demonstrated the regulation of *KCNN1* by these fusion proteins, which bind at GGAA microsatellites near *KCNN1* promoter. In addition, *KCNN1* has been shown to be involved in the regulation of ES cell proliferation, its silencing being associated with a slowing of the cell cycle. Finally, *KCNN1* expression modulates membrane potential and calcium flux suggesting the role of calcium in *KCNN1* driving cell proliferation. These results highlight that *KCNN1* is a direct EWS-FLI1 and EWS-ERG target, and is involved in the regulation of ES cell proliferation, making it an interesting therapeutic target in ES.

## Introduction

With an incidence of 1 case per 1.5 million population (Riggi *et al*, 2021), Ewing sarcoma (ES) is the second most common primary malignant bone tumor in children, adolescents and young adults. At the genetic level, ES is characterized by a chromosomal translocation between a member of the FET family and a member of the ETS family (Grünewald *et al*, 2018). In 85% of cases, the chromosomal translocation found is (11;22)(q24;q12), between the EWS RNA-binding protein and the FLI1 transcription factor, leading to the EWS-FLI1 fusion protein (Delattre *et al*, 1992). This chimeric protein acts as an oncogene factor for various transcription factors (Cidre-Aranaz & Alonso, 2015), such as Gli1 (Mullard *et al*, 2020; Merchant *et al*, 2009), c-Myc (Dauphinot *et al*, 2001) or DAX1 (Kinsey *et al*, 2006; Mendiola *et al*, 2006). In 10% of cases, the chromosomal translocation leads to the emergence of the EWS-ERG chimeric protein (Zucman *et al*, 1993). Current treatment consists of neoadjuvant and adjuvant chemotherapy (including vincristine, ifosfamide, doxorubicin and etoposide) (Strauss *et al*, 2021), and surgical resection of the tumor. This increases patient survival to 70%, which unfortunately decreases to less than 30% when patients are resistant to chemotherapy or when lung metastases are present at diagnosis. The lack of improvement in these survival rates over the past decades indicates the urgent need for new therapies.

In recent years, studies have emerged on the involvement of ion channels in tumorigenesis and the usefulness of targeting them in a cancer treatment strategy (Peruzzo & Szabo, 2019; Catacuzzeno *et al*, 2021; Potier-Cartereau *et al*, 2022a; Prevarskaya *et al*, 2018). For example, some calcium or potassium channels have been shown to be involved in gastrointestinal cancer (Anderson *et al*, 2019), colorectal cancer (Ibrahim *et al*, 2019) or myeloma (Wang *et al*, 2014). Among these channels, potassium channels form the largest family, with no less than 70 genes encoding these structures. They are subdivided into 4 classes, according to the number of transmembrane segments and pore domains (Girault *et al*, 2012). Among these 4 classes, calcium-activated potassium channels (KCa) are divided into 3 subclasses based on their unitary conductance: small conductance (SKCa: SK1, SK2 and SK3), intermediate conductance (IKCa: SK4) and big conductance (BKCa: BK) (Girault *et al*, 2012). SKCa channels active form consists of a homo-heterotetramer of α subunits, each consisting of 6 transmembrane segments and a pore domain. SK3, whose α subunit is encoded by the *KCNN3* gene, has previously been shown to be involved in bone metastasis from breast cancer (Potier *et al*, 2006), and overexpressed in prostate cancer (Bery *et al*, 2020, 2021) and melanoma (Chantome *et al*, 2009; Tajima *et al*, 2006), just like *KCNN2*, encoding SK2 channel (Tajima *et al*, 2006), in addition to pancreas cancer (Rapetti-Mauss *et al*, 2023).

While the involvement of *KCNN1* expression has never been described in a key biological function of cancer development, this study shows for the first time the role of *KCNN1* in the tumorigenesis of ES. We demonstrated that (i) among all tumors studied, *KCNN1* is exclusively expressed in biopsies from ES patients and in ES cell lines, (ii) *KCNN1* expression is not found significant in any healthy tissue except the brain, (iii) among the 8 *KCNN1* transcripts described in the literature, 3 are predominantly expressed in ES cells, (iv) EWS-FLI1 regulation of these 3 *KCNN1* transcripts, expressed in ES cells, is performed through binding in open chromatin region enriched with GGAA sequence motif, (v) the EWS-ERG fusion protein is able to regulate *KCNN1* expression by a mechanism similar to EWS-FLI1, (vi) *KCNN1* expression level is involved in the regulation of ES cell proliferation and the ability of ES cells to form colonies, (vii) *KCNN1* gene expression controls plasma membrane potential and calcium influx through Store-Operated Calcium Entry (SOCE) and Constitutive Calcium Entries (CCE) by increasing driving force for calcium, and (vii) *KCNN1* expression is inversely correlated with patient survival.

## Results

### *KCNN1* is overexpressed in Ewing sarcoma patients and cell lines

To address the potential role of SKCa channels in ES development, the expression of genes encoding the different SKCa channels (SK1, SK2 and SK3) was first compared in ES biopsies, using the public R2 available databases (https://r2.amc.nl). RNA-seq data from a cohort of 117 ES patients show that *KCNN1* (encoding SK1) is significantly more expressed in ES patients than *KCNN2* or *KCNN3,* encoding respectively SK2 and SK3 (Fig 1A). Comparison of *KCNN1* expression across multiple tumors’ biopsies shows that *KCNN1* is exclusively highly expressed in biopsies from ES patients (Fig 1B) unlike *KCNN2* and *KCNN3,* expressed in various tumors (Fig EV1). *KCNN2* is indeed highly expressed in glioma, glioblastoma and prostate biopsies (Fig EV1A), while *KCNN3* is ubiquitously expressed in biopsies of various tumors (Fig EV1B). Interestingly, according to the Kaplan Meier curve, *KCNN1* expression is significantly inversely correlated with patient survival at 5 years after diagnosis (Fig 1C). Indeed, the probability of overall survival at 72 months is almost divided by 3 when patients strongly expressed *KCNN1*, compared to patients presenting low expression of *KCNN1* (Fig 1C). Regarding *KCNN1* expression in healthy tissues, analysis across RNA-Seq samples from Genotype-Tissue Expression (GTEX, normal tissue, Fig 1D) and in bone marrow-Mesenchymal Stem Cells (MSCs, R2 database), the putative cell of origin of ES (Fig 1E), shows higher expression of *KCNN1* only in brain tissues. Other healthy tissues (Fig 1D) or MSCs (Fig 1E) barely express *KCNN1*.

**Figure 1:**
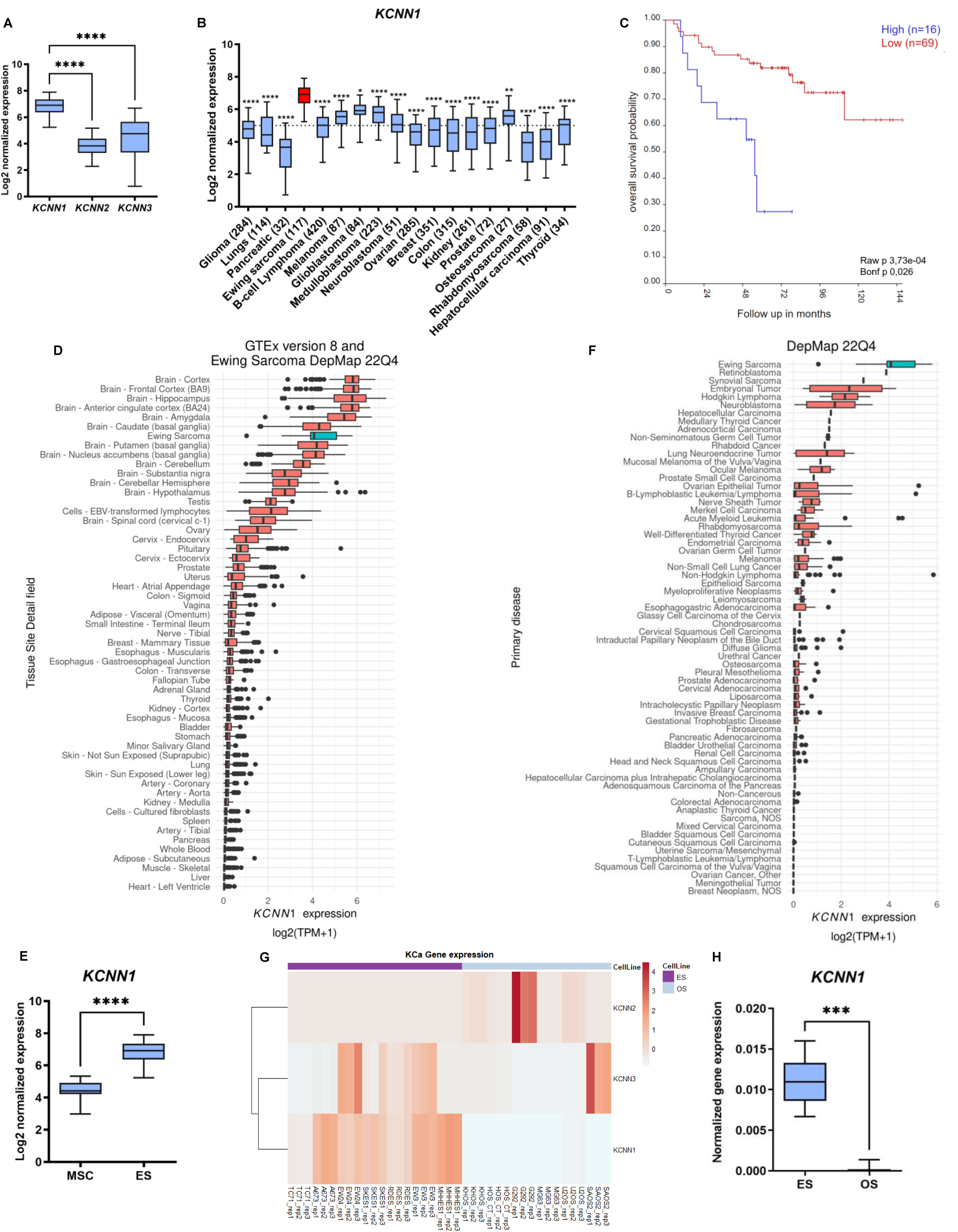
*KCNN1* is overexpressed in biopsies from ES patients and in ES cell lines. A Gene expression (Log2 normalized expression) of genes encoding small conductance calcium-activated potassium channels (*KCNN1, KCNN2* and *KCNN3*) in Ewing sarcoma patients (117 patients) (Brown-Forsythe and Welch Anova test). B Gene expression (Log2 normalized expression) of *KCNN1* in biopsies originated from patients with different tumors (Kruskal-Wallis test). C Overall survival probability (Kaplan Meier curve) of patients suffering from Ewing sarcoma depending on *KCNN1* expression (high or low). The Bonf p corresponds to the p-value after the Bonferroni correction for multiple comparisons. D Expression of *KCNN1* in normal tissues. TPM, transcripts per million. E Gene expression (Log2 normalized expression) of *KCNN1* in Mesenchymal Stem Cells (MSCs) coming from thirty healthy donors (Yamaguchi database – 30 donors) and in patients suffering from Ewing sarcoma (ES, Olivier Delattre database - 117 patients) (Unpaired t-test). F Expression of *KCNN1* in normal tissues. G Heat-map showing color-coded expression of small conductance calcium-activated potassium channels (SKCa) in seven Ewing sarcoma cell lines (ES) and six osteosarcoma cell lines (OS). Counts were variance stabilized using DESeq2’s rlog transformation function and transformed to z-score. H Validation of *KCNN1* normalized gene expression in Ewing sarcoma cell lines and osteosarcoma cell lines. (N=7). The boxes indicate the first and third quartiles - and the midline represents the median - of the normalized *KCNN1* mRNA expression determined by RT-qPCR. The whiskers indicate the fifth and the ninety-fifth percentiles (Mann-Whitney test). Data information: In (A-B, E and H), the boxes boundaries indicate the first and third quartiles - and the midline represents the median – of genes expression in Log2. The whiskers indicate the fifth and the ninety-fifth percentiles. In (D and F), boxes represent the interquartile range, upper and lower whiskers the largest and smallest values respectively. TPM: Transcript Per Million. *: p < 0,05 ; **: p< 0,01 ; ***: p<0,001 and ****: p<0,0001.

To validate these findings at the cellular level, *KCNN1* expression was assessed in 67 different tumor-derived cell lines (Fig 1F). *KCNN1* shows restricted expression in ES and retinoblastoma (Fig 1F). It is important to note that *KCNN1* expression in retinoblastoma was performed in only one cell line in contrast to the evaluation of *KCNN1* expression in ES, which was performed in 18 samples. These results were confirmed by comparing *KCNN1*, *KCNN2*, and *KCNN3* expression in 7 ES and in 6 osteosarcoma (OS) cell lines (GSE229906, Fig 1G). *KCNN1* is exclusively expressed in ES cell lines, as confirmed by RT-qPCR (Fig 1H), in contrast to *KCNN2* and *KCNN3* expressed in OS, or in both ES and OS cell lines, respectively (Fig 1G).

Taken together, these results show that i) *KCNN1* is exclusively overexpressed in biopsies from ES patients and in ES cells, and ii) that *KCNN1* expression is associated with patient survival at 5 years after diagnosis.

### *KCNN1* is transcriptionally regulated by the EWS-FLI1 fusion protein

Since EWS-FLI1 is present in 85% of ES cases, the regulation of *KCNN1* expression by this fusion protein was then investigated, using a function extinction approach. For this purpose, A673/TR/shEF cells, an ES cell line stably transduced with a doxycycline-induced shRNA targeting *EWS-FLI1*, referred to as ASP14 in this article, were used (Carrillo *et al*, 2007). Doxycycline treatment of ASP14 cells induces a progressive decrease in *EWS-FLI1* expression after 4 hours of doxycycline treatment, until reaching 95% silencing after 24 hours of doxycycline treatment (Fig 2A, upper panel). Following a similar trend, *KCNN1* expression is decreased as early as 14 hours of cell treatment and this decrease is maintained up to 24 hours of treatment (Fig 2A, lower panel). To confirm these results in other ES cell lines, a pool of siRNAs targeting *EWS-FLI1* was used in A673, RDES and SKES1 cell lines (Fig 2B). In these three cell lines, silencing of *EWS-FLI1* expression by siRNAs (Fig 2B, left panels for each cell line) leads to a decrease in *KCNN1* expression of 75%, 40% and 50% in A673, RDES and SKES1, respectively (Fig 2B, right panels for each cell line).

**Figure 2:**
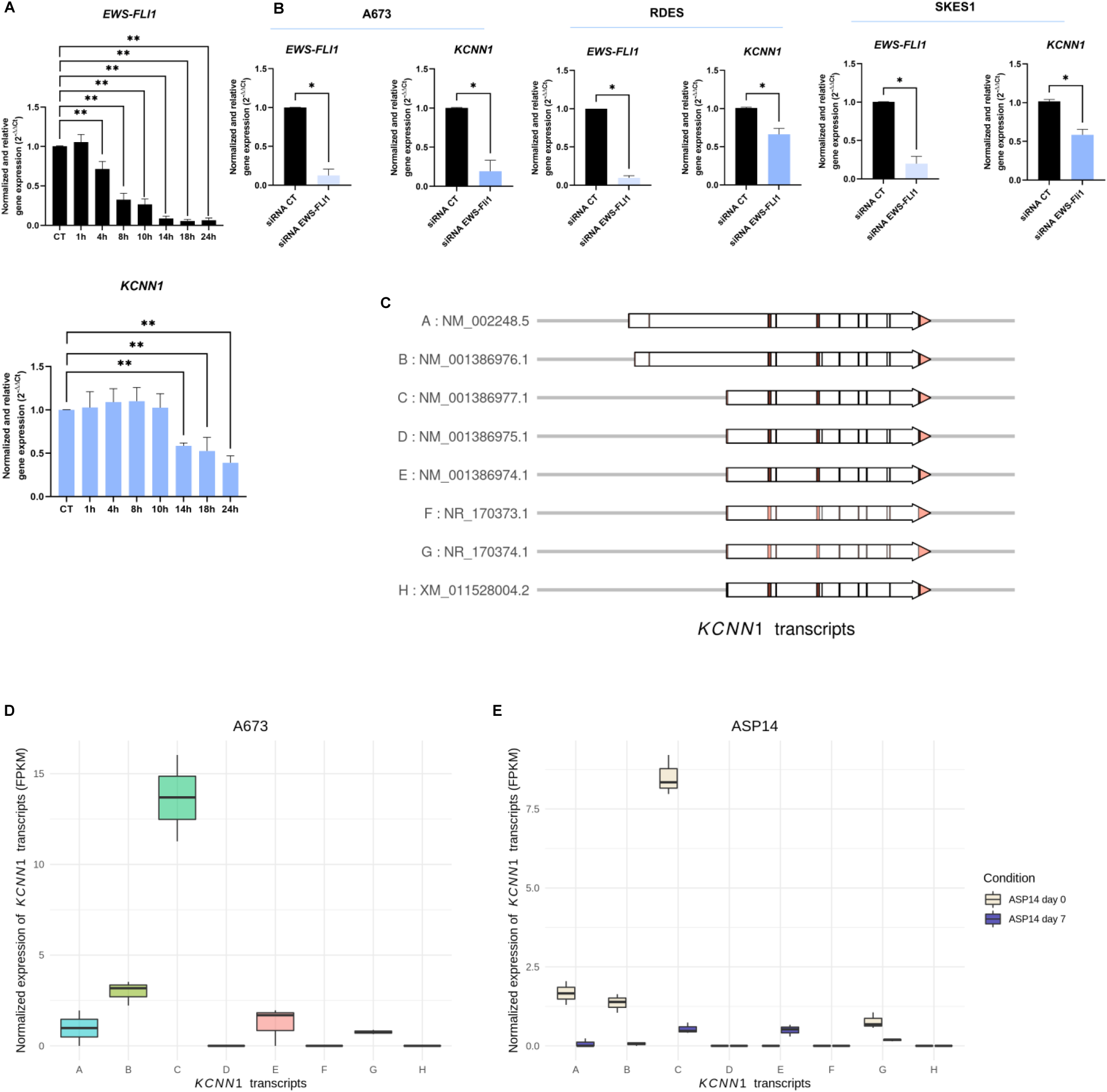
*KCNN1* is regulated by the chimeric protein EWS-FLI1. A Normalized and relative *EWS-FLI1* (upper figure) and *KCNN1* (lower figure) gene expressions in ASP14 cell line after doxycycline treatment (1 µg/mL) (N=5). B Normalized and relative *EWS-FLI1* (left panel for each cell line) and *KCNN1* (right panel for each cell line) gene expressions in A673 (N=3), RDES (N=4), and SKES1 (N=3) cell lines after 48h of transitory transfection with a pool of siRNA targeting *EWS-FLI1* (siRNA EWS-FLI1) or a siRNA control (siRNA CT). C Different *KCNN1* transcripts according to their position on the genome. Vertical bars represent exons. Light pink variants are non-coding transcripts. D Normalized expression of *KCNN1* transcripts (FPKM) in A673 cell line (N=3). E Normalized expression of *KCNN1* transcripts (FPKM) in ASP14 cell line at day 0 and day 7 of doxycycline treatment (1 µg/mL). Data information: In (A-B), Bars indicate means ± SD of relative and normalized *EWS-FLI1or KCNN1* mRNA expression determined by RT-qPCR. In (D-E), FPKM : Fragment Per Kilobase Per Million reads. *: p < 0,05 ; **: p< 0,01 (Mann-Whitney tests).

Among the 8 *KCNN1* transcripts described in the literature (NCBI RefSeq) (Fig 2C), analysis of RNA-Seq data based on an alignment on the transcriptome sequence shows that A673 cells express the 4 transcripts A (NM_002248.5), B (NM_001386976.1), C (NM_001386977.1) and E (NM_001386974.1). The other 6 ES lines studied significantly expressed only the transcripts, A, B and C (Fig EV2A). It is notable that the transcript C is predominantly expressed in ES cells (Fig 2D), and that, based on the predictions of the SK1 channel structures of the different variants A, B and C (Fig EV2B), the C transcript generates a subunit with a deletion of 3 amino acids in the calmodulin binding domain, compared to the one obtained with the canonical transcript (i.e the A transcript). However, as it can be seen, there is no drastic change observed in the structure of the SK1 channel encoded by the *KCNN1* C transcript, compared with the channel encoded by the *KCNN1* A transcript (Fig EV2B). Interestingly, *EWS-FLI1* silencing in ASP14 cells results in a decrease of the expression level of the 3 major *KCNN1* transcripts in ES cells (GSE164373, (Buchou *et al*, 2022; Data ref: Buchou *et al*, 2022)) (Fig 2E). In particular, the expression of the highly expressed C transcript is drastically decreased after 7 days of treatment of the cells with doxycycline. The lower expressed transcripts A and B also have decreased expression (Fig 2E).

ChIP-Seq data (GSE176400, (Orth *et al*, 2022; Data ref: Orth *et al*, 2022)) on the A673 cell line (Fig 3A, left panel) show a significant peak of H3K4me3 (a histone modification marking active promoters (Rada-Iglesias *et al*, 2011)) at the promoter of *KCNN1* C transcripts, as well as a significant peak of H3K27ac (a histone modification marking active promoters and enhancers (Wang *et al*, 2008)) and a FLI1 peak (with significant q. value) in close proximity to the C transcript promoter. Our results seem to correspond to the positions observed by Fuest and colleagues, concerning the binding of EWS-FLI1 to an alternative enhancer in the third intron of *KCNN1* A transcript (Fuest *et al*, 2022). The same results can be observed for the 11 other ES cell lines characterized by the EWS-FLI1 fusion protein (Fig EV3A). Interestingly, a similar regulatory mechanism by the EWS-ERG fusion protein can be observed for EW3, CHLA25, and TC106, 3 cell lines characterized by the presence of the EWS-ERG fusion protein (Fig 3A, middle and right panels). These EWS-FLI1/EWS-ERG binding sites contain at least 12 GGAA microsatellites repetitions, indicated by the yellow vertical lines (Fig 3A-D).

**Figure 3:**
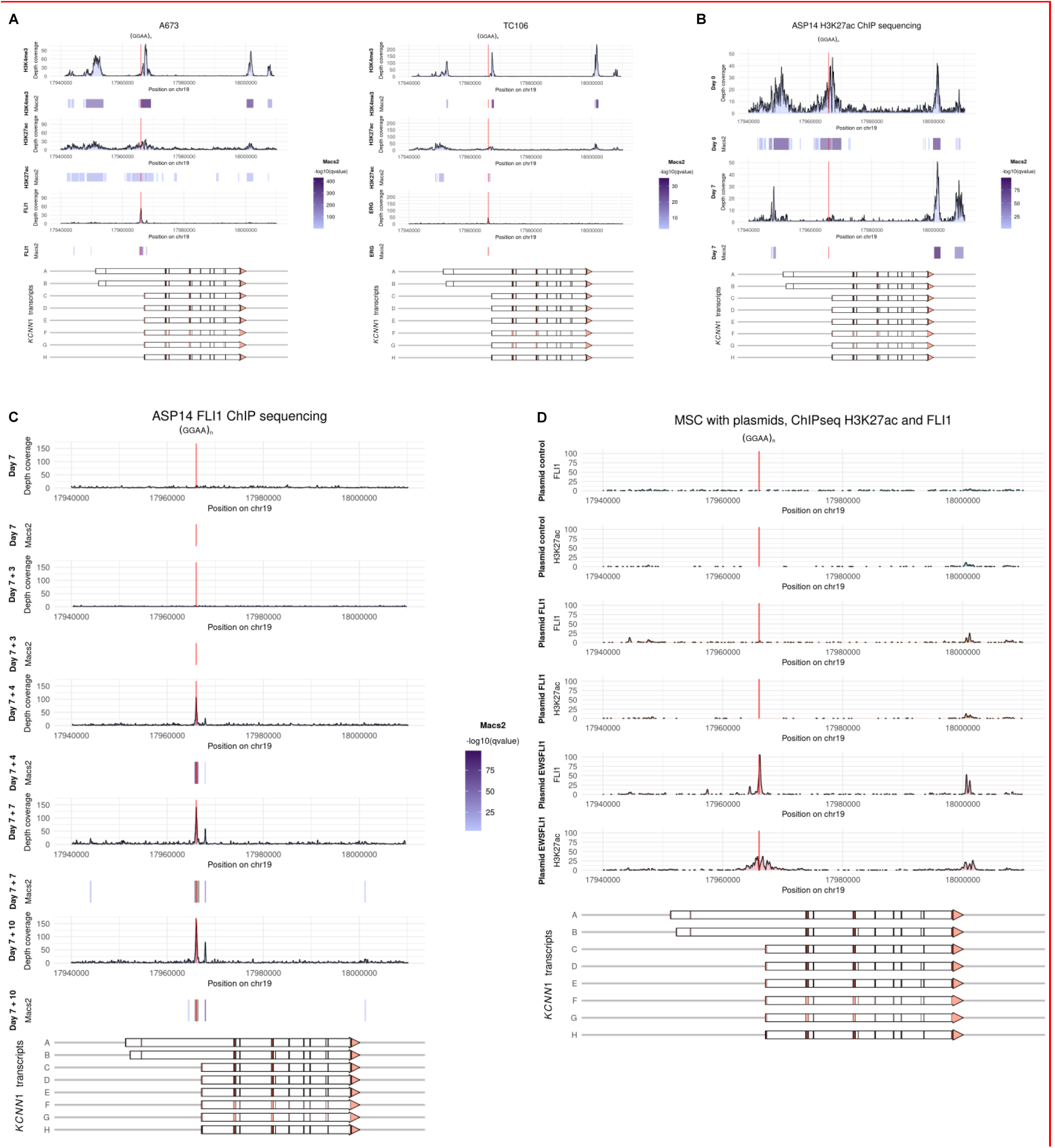
EWS-FLI1 regulates *KCNN1* expression by binding GGAA microsatellites. A H3K4me3, H3K27ac and FLI1 ChIP-seq data with quantitative value of peaks around *KCNN1* promoters and sequences of EWS-FLI1 binding sites in A673 and TC106 cell lines. B H3K27ac ChIP-Seq of ASP14 cell line at day 0 and day 7 of doxycycline treatment with quantitative value of peaks around *KCNN1* promoters. C FLI1 ChIP-Seq of ASP14 cell line with quantitative value of FLI1 peaks around *KCNN1* promoters, at day 7 of doxycycline treatment, and 3 days (day 10), 4 days (day 11), 7 days (day 14) and 10 days (day 17) after discontinuation of doxycycline treatment. D H3K27ac and FLI1 ChIP-Seq data on MSCs transfected with control plasmid, FLI1 overexpression plasmid or EWS-FLI1 overexpression plasmid around *KCNN1* promoters.

To confirm the binding of EWS-FLI1 around *KCNN1* promoter regions of the A, B and C transcripts, publicly available H3K27ac (Fig 3B) and FLI1 (Fig 3C) ChIP-Seq data performed on the ASP14 cell line (GSE129155, (Aynaud *et al*, 2020; Data ref: Aynaud *et al*, 2020)) were used. Two significantly called H3K27ac peaks are observed around *KCNN1* promoter domain in the absence of doxycycline treatment (day 0), indicating areas of open chromatin in these regions of the *KCNN1* A, B, and C promoter transcripts (Fig 3B, upper panel). Interestingly, the amplitude of the peak near *KCNN1* C transcript decreases after 7 days of treatment, i.e after silencing of EWS-FLI1 (Fig 3C), which confirms the decreased expression of *KCNN1* shown in Figure 2A. FLI1 ChIP-Seq data show EWS-FLI1 binding at this open chromatin region (Fig 3C), with a FLI1 peak associated with the H3K27ac activation mark (Fig 3B). Indeed, FLI1 peaks gradually reappear at the chromatin-opening region after re-expression of EWS-FLI1 following the interruption of treatment with doxycycline (Fig 3C). Note that FLI1 peaks are specific to the EWS-FLI1 fusion protein since the FLI protein is absent in ES cells due to chromosomal translocation (Franzetti *et al*, 2017). It can be noted that EWS-FLI1 silencing leads to an increase in *KCNN3* expression (Fig EV3B) with EWS-FLI1 binding being observed in a chromatin-opening area in the vicinity of the *KCNN3* promoter (Fig EV3C), the intensity of the peak being however less important than for *KCNN1* (Fig 3C). Following a shRNA-mediated silencing approach in ES cells, overexpression of EWS-FLI1 in the putative cell of origin of ES, MSCs, was used (GSE94278, (Boulay *et al*, 2017; Data ref: Boulay *et al*, 2017)). Neither H3K27ac nor FLI1 peak is observed around *KCNN1* promoter areas in MSCs transfected with the control plasmid (Fig 3D, upper panel) or FLI1 overexpression plasmid (Fig 3D, medium panel). In contrast, when EWS-FLI1 was overexpressed, H3K27ac and EWS-FLI1 peaks appear around the promoter domain of *KCNN1* transcript C (Fig 3D, lower panel).

Altogether, these results show transcriptional regulation of *KCNN1* by EWS-FLI1 or EWS-ERG, enabled by their binding to GGAA microsatellites in chromatin opening regions.

### *KCNN1* is involved in the regulation of ES cell proliferation

To pursue the investigation on the potential role of SK1 channel in ES development, the involvement of *KCNN1* in the regulation of ES cell cycle was studied. To examine this hypothesis, doxycycline-inducible shRNAs targeting *KCNN1* were used. The efficacy of shRNA was first evaluated on *KCNN1* expression after 72 hours (Fig 4A and Fig EV4A) and 7 days (Fig EV4B) of treatment of A673 with doxycycline, respectively. After 72 hours of doxycycline treatment, a decrease of at least 70% of *KCNN1* expression can be observed in all 3 shRNA targeting *KCNN1* cells (i.e shRNA 511, shRNA 908 and shRNA 576 cells), compared to the shRNA control (shRNA CT) cells (Fig 4A). In addition, RNA-Seq analysis indicates that the transcript mainly expressed in A673 shRNA 908 cells, the C transcript, has its expression reduced after treatment of the cells with doxycycline (GSE229906, Fig EV4A). And, after 7 days, there is still a decrease by 65% of *KCNN1* expression in all 3 shRNA cells (Fig EV4B).

**Figure 4:**
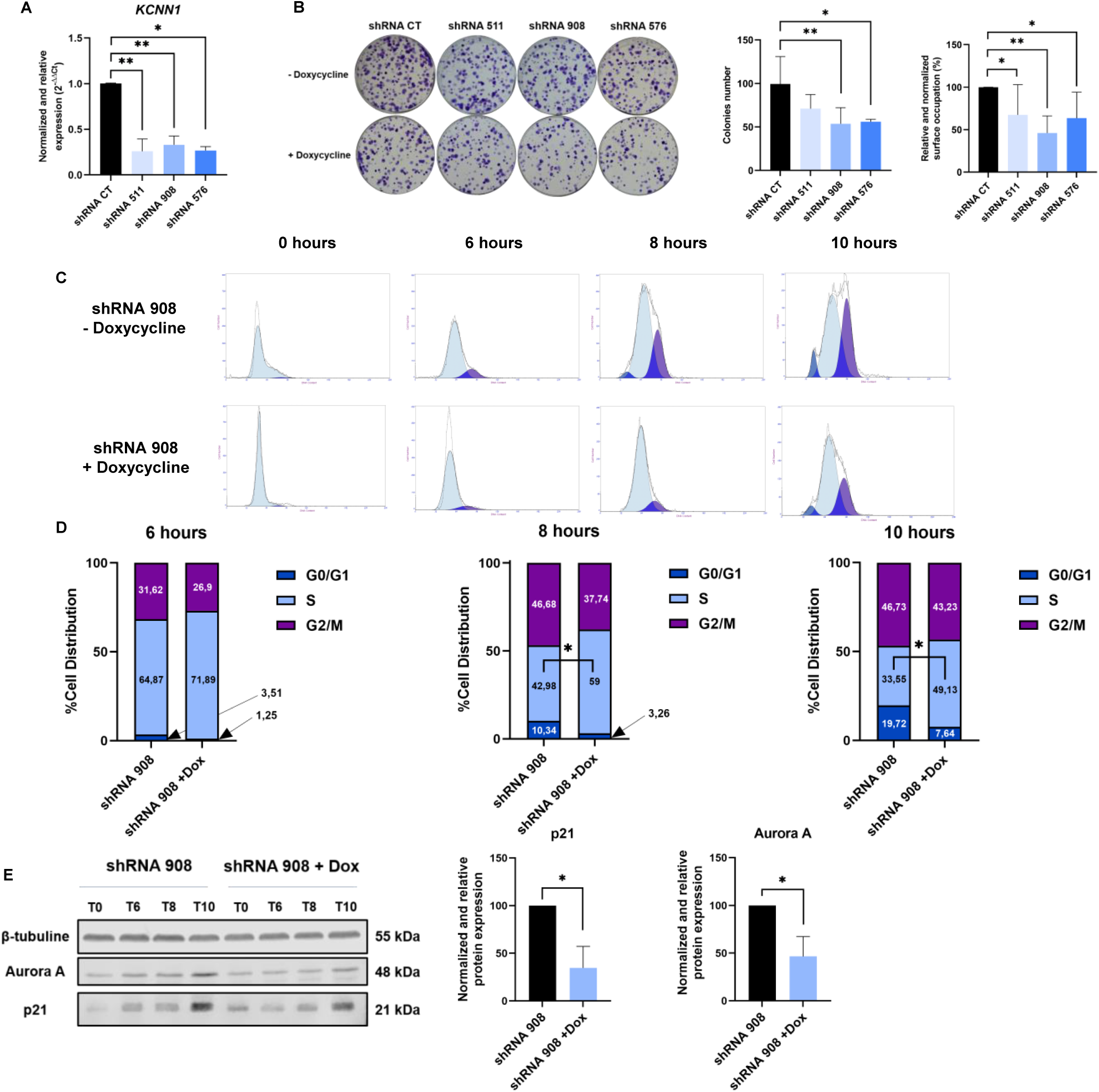
*KCNN1* is involved in the regulation of ES cell proliferation. A Verification of *KCNN1* extinction in A673 cell line stably transduced with doxycycline-inducible shRNA control cells (shRNA CT) or 3 shRNA cells targeting *KCNN1* (shRNA 511, shRNA 908 and shRNA 576) after 72 hours of treatment with doxycycline (0,5 µg/mL). Bars indicate mean ± SD of relative and normalized *KCNN1* mRNA expression obtained after RT-qPCR (N=5 for shRNA CT and shRNA 908, N=4 for shRNA 511 and N=3 for shRNA 576). B Left panel: clonogenicity assays of A673 cell line stably transduced with shRNA control (shRNA CT) or 3 shRNA targeting *KCNN1* cells (shRNA 511, shRNA 908 and shRNA 576) after 72 hours of doxycycline treatment (0,5 µg/mL). Each colony was dyed in crystal violet after being fixed with glutaraldehyde. Medium panel: Colonies numbers observed on the left panel. Bars indicate mean ± SD of counted colonies on the left panel. Right panel: Relative surface area of clones observed on the left panel. Bars indicate mean ± SD of the clones surface area observed on the left panel relative to the cells not treated with doxycycline (N=6 for shRNA CT, N=3 for shRNA 511 and shRNA 908 and N=7 for shRNA 908). C Cell cycle assays on shRNA 908 cells +/-doxycycline 0 hours, 6 hours, 8 hours or 10 hours after thymidine double block. Dark blue: G0/G1 phase, Light blue: S phase, Purple: G2/M phase. D Percentage of cell distribution 6 hours, 8 hours and 10 hours after thymidine double block (N=5 for 6 and 10 hours after thymidine double block ; N=6 for 8 hours after thymidine double block, Wilcoxon test). E Western blots of p21 and Aurora A (left panel) and their quantification normalized to β-tubulin 10 hours after the thymidine double block (medium and right panel) (N=3). Data information: *: p < 0,05 ; **: p< 0,01 (A-B: Mann-Whitney tests).

To investigate the involvement of *KCNN1* in the regulation of cell proliferation, a key function during tumor development, clonogenicity assays were performed. Seventy-two hours after treatment of the cells transduced with 3 shRNA targeting *KCNN1* with doxycycline, a decrease of the ability of shRNA cells to form colonies is observed (Fig 4B). Indeed, a significant decrease in the number of clones is observed with a decrease of at least 50% for shRNA 908 and shRNA 576 cells (Fig 4B, middle panel). Clones size is also significantly smaller for all 3 shRNA cells, up to 50% for shRNA 908, compared to shRNA CT (Fig 4B, right panel). These results were confirmed with viability assays on MHHES1 cells transfected with a pool of siRNAs targeting *KCNN1* (Fig EV4C). A slowdown of cell proliferation is indeed observed during the time lapse, 48 hours after transfection. To better understand the involvement of *KCNN1* in the regulation of ES cell proliferation, cell cycle assays on synchronized cells (with thymidine double block) were then realized (Fig 4C-D). On average, 95% of cells are synchronized in S phase at the end of the double block (Fig 4C). Six hours after the end of the double block, more cells are in the G2/M phase in the control condition (31,62%), compared to *KCNN1* knocked-down cells (26,9%). This slowdown of the cell cycle is even more pronounced 8 hours and 10 hours after the end of the double block, the majority of shRNA 908 cells treated with doxycycline still being in the S phase (49,13% vs 33,55%). In addition, more control cells are in the G0/G1 phase 10 hours after the thymidine double block (19,72%) than after *KCNN1* extinction (7,64%).

Measurement of p21 and Aurora A expression, known to accumulate in G2/M phase, confirms the slowing down of the cell cycle after *KCNN1* expression silencing (Fig 4E). Indeed, 10 h after stopping the blockade of cells in S phase by thymidine treatment, a higher expression of p21 and Aurora A is measured, illustrating a faster return of cells to G2/M phase in control condition than after silencing of *KCNN1* expression (Fig 4E, medium panels and Fig EV4D).

Together, these results show the involvement of *KCNN1* expression in the regulation of ES cell cycle and the ability of cells to form colonies.

### *KCNN1* expression modulates calcium flux

As it is known that SKCa controls membrane potentials of various cells including cancer cells, firstly, *KCNN1* involvement in membrane potential regulation was investigated. Functionally, *KCNN1* silencing leads to plasma membrane depolarization in a 2D or 3D cell model (Fig 5A and 5B). Indeed, the DiBAC4(3) fluorescence measurement is significantly higher in shRNA 511, shRNA 908 and shRNA 576 cells than in shRNA CT cells (Fig 5A). In addition, measured membrane potentials using intracellular microelectrodes on shRNA 908 spheroids shows a significant depolarization of 20 mV compared to the shRNA CT spheroids (Fig 5B right panel). As expected, DiBAC4(3) fluorescence measurements showed the ability of an activator of SK1, compound GW542573X, to hyperpolarize the cell membrane (Fig 5C).

**Figure 5:**
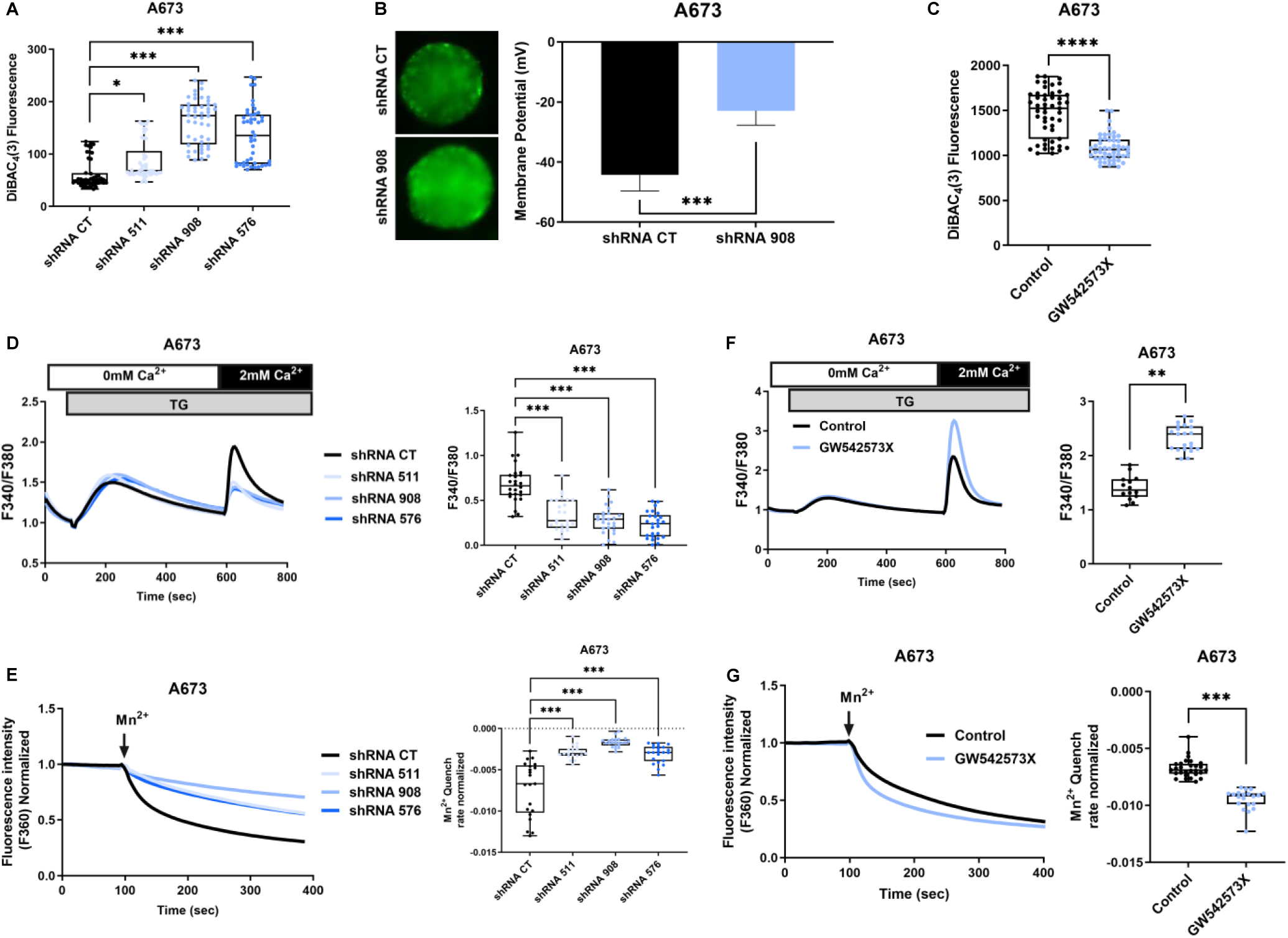
*KCNN1* expression modulates membrane potential and calcium flux. A Measures of the plasma membrane depolarization in A673 cell line stably transduced with shRNA control cells (shRNA CT) or 3 shRNA cells targeting *KCNN1* (shRNA 511, shRNA 908 and shRNA 576) after 7 days of doxycycline treatment (0,5 µg/mL). The boxes indicate the first and third quartiles - and the median inside - of the normalized DiBAC_4_(3) fluorescence. (N=48 for shRNA CT and shRNA 908, N=46 for shRNA 511, N=45 for shRNA 576). B Images (left panel) and measures of membrane potential (right panel) of shRNA CT or shRNA 908 spheroids treated 7 days with doxycycline (0,5 µg/mL) (N=7). C Measures of the plasma membrane depolarization in A673 cell line treated with SK1 activator, GW542573X (N=50). D Effect of *KCNN1* knockdown on SOCE in A673 cell line stably transduced with shRNA control (shRNA CT) or 3 shRNA targeting *KCNN1* (shRNA 511, shRNA 908 and shRNA 576). Left panel: Fluorescence measurement of SOCE induced by thapsigargin (TG). Right panel: Fluorescence measurement of SOCE induced by TG in shRNA *KCNN1* conditions compared to control on the relative fluorescence of shRNA CT cells loaded with Fura2-AM (N=26) (Unpaired t-test). E Effect of *KCNN1* knockdown on constitutive calcium entries using Mn2+ quenching assay. Left panel: Effect of *KCNN1* knockdown on Mn2+ quenching rate. Right panel: Effect of *KCNN1* knockdown on normalized Mn2+ quenching slope (N=21 for shRNA CT, N=20 for shRNA 511 and N=22 for shRNA 908 and shRNA 576). F Effect of SK1 activator, GW542573X, on SOCE in A673 cell line. Left panel: Fluorescence measurement of SOCE induced by TG. Right panel: Fluorescence measurement of SOCE induced by TG in GW542573X conditions compared to control on the relative fluorescence of shRNA CT cells loaded with Fura2-AM (N=14 for Control and N=21 for G542573X condition). G Effect of SK1 activator GW542573X, on constitutive calcium entries using Mn2+ quenching assay. Left panel: Effect of the GW542573X on Mn2+ quenching rate. Right panel: Effect of the GW542573X on normalized Mn2+ quenching slope (N=28 for Control and N=20 for GW542573X condition). Data information: In (A and C), the boxes indicate the first and third quartiles - and the median inside - of the normalized DiBAC_4_(3) fluorescence. The whiskers indicate the fifth and the ninety-fifth percentiles and the points, each replicate. In (D and F) the boxes indicate the first and third quartiles - and the median inside - of the F340/F380 ratio. The whiskers indicate the fifth and the ninety-fifth percentiles and the points, each replicate. In (E and G), the boxes indicate the first and third quartiles - and the median inside - of the normalized Mn^2+^ Quench rate. The whiskers indicate the fifth and the ninety-fifth percentiles and the points, each replicate. *: p < 0,05 ; **: p< 0,01 ; ***: p<0,001 (A and D-E: Anova tests ; B and F-G : Mann-Whitney tests).

Then, to identify the molecular mechanisms by which the *KCNN1* expression affect ES cell proliferation, the effect of its silencing on calcium signaling was investigated. Store-Operative Calcium Entry (SOCE) and Constitutive Calcium Entries (CCE) were measured using Fura-2AM and the Mn^2+^ quenching technique, respectively. Under doxycycline treatment (i.e after silencing of *KCNN1* expression), there is a significant decrease of calcium entry after SOCE activation by thapsigargin (TG), following addition of 2 mM calcium in extracellular solution, in *KCNN1* knocked-down cells in comparison to the shRNA CT cells (Fig 5D). Silencing *KCNN1* expression after treatment of the cells with doxycycline also reduces Mn^2+^ quenching rate (Fig 5E) in all 3 shRNA cells, compared to the shRNA CT cells, indicating decreased CCE. On the other hand, using the SK1 channel activator GW542573X, an increase of both SOCE and CCE is observed in the A673 cell line (Fig 5F-G). The decrease of calcium entries is also observed in *KCNN1* knocked-down SKES1 cells compared with the CT shRNA cells (Fig EV5A-B).

Overall, these results indicate that *KCNN1* expression is involved in the control of membrane potential and calcium influx in ES cells.

## Discussion

Despite advances in patient management and improved clinical outcomes for patients with Ewing sarcoma (ES), anti-ES treatment remains challenging and there is an urgent need for new therapeutic strategies. Intercellular communication is essential for maintaining tissue homeostasis, and this exchange is mediated by membrane structures known as ion channels. Among these channels, the potassium channel family is the most important with in particular calcium-activated potassium channels (KCa) including those with small conductances (SKCa) (Kuang *et al*, 2015). Abnormal expression and dysfunction of ion channels can give rise to various cancers (Prevarskaya *et al*, 2010; Comes *et al*, 2015). In the last few years, potassium channels such as KCa have thus emerged as key molecular targets for the development of cancer treatments (Prevarskaya *et al*, 2018; Bates, 2015; Huang & Jan, 2014; Pardo & Stühmer, 2014; Capatina *et al*, 2020; Ohya *et al*, 2021). KCa channels have indeed been implicated in the tumorigenesis process in patients and in various animal models (Ibrahim *et al*, 2019; Tajima *et al*, 2006; Mohr *et al*, 2019). In particular, among this subfamily, the small conductance SK3 channel plays a major role in the control of cell migration (Potier *et al*, 2006) and thus in the development of bone metastases (Chantôme *et al*, 2013). Our work describes for the first time to our knowledge, the expression of the *KCNN1*, gene encoding SK1, in a cancer tissue, i.e ES. Indeed, our results on biopsies from patients with different cancers show an expression of *KCNN1* exclusively in ES. Although Fuest *and al*., (Fuest *et al*, 2022) did not observe any correlation between the 16-year survival rate and *KCNN1* expression, our results showed a significant inverse correlation between *KCNN1* expression and the 5-year survival rate. Looking at survival rates at different times and the different technics of analyses might explain these results, but the same tendency being observed, our work highlights the importance of *KCNN1* expression in patient survival.

In this context, we can hypothesize that *KCNN1* could be a specific biomarker for ES. In immunohistochemistry, ES cells are characterized by intense membrane labeling with the anti-CD99 antibody, directed against the *MIC2* gene product (Ambros *et al*, 1991). However, this sensitive staining is not very specific, as it is found in other sarcomas such as synovialosarcoma, desmoplastic tumor, mesenchymal chondrosarcoma and some lymphoma and lymphoblastic leukemia (Baldauf *et al*, 2018). The anti-FLI1 antibody is neither sensitive nor specific for ES (Mhawech-Fauceglia *et al*, 2007) and immunostaining with the anti-NKX2.2 antibody, a transcription factor involved in glial/neuroendocrine differentiation, is reported to be very sensitive, but lacks specificity for ES, which limits its diagnostic utility (Hung *et al*, 2016). A signature of ES represented by a specific and sensitive biomarker appears to thus be an ideal tool to detect tumors early and improve prognosis through early management. Nevertheless, the presence of SK1 channel at the membrane, not observed by Fuest *and al*., (Fuest *et al*, 2022), and the development of immunostaining with a specific antibody remains to be done. Furthermore, our work shows very low expression of *KCNN1* in healthy tissues, with the exception of the brain, complementing and reinforcing the study of Fuest and colleagues, who demonstrated specificity of *KCNN1* expression in ES cell lines compared to healthy bone tissue (Fuest *et al*, 2022). It can therefore be hypothesized that targeting *KCNN1* or a potential SK1 channel with a strategy using a candidate that does not cross the blood-brain barrier would limit the development of side effects.

Several studies have shown that EWS-FLI1 is able to regulate various genes expression (Cidre-Aranaz & Alonso, 2015), such as *Gli1* (Mullard *et al*, 2020; Merchant *et al*, 2009), or more recently *LOXHD1* (Deng *et al*, 2022). Both FLI1 and the fusion protein EWS-FLI1 bind the canonical ETS binding motif, GGAA (Wasylyk *et al*, 1993; Boeva *et al*, 2010). It has also been proved that EWS-FLI1 is able to bind GGAA microsatellites (Gangwal *et al*, 2008; Guillon *et al*, 2009), which can be repeated up to 20 times. This fusion protein is known to present the ability to recruit chromatin remodeling complexes and the transcription machinery (Boulay *et al*, 2017), which can lead to the emergence of variant transcripts (Deng *et al*, 2022). Here, we firstly observed that *KCNN1* expression is regulated by EWS-FLI1, confirming the previous study of Fuest and colleagues (Fuest *et al*, 2022). Indeed, EWS-FLI1 binds GGAA microsatellites close to the promoter of *KCNN1* shorter transcripts. In this context, it has been shown in previous studies that EWS-FLI1 might be able to change the 3D configuration of chromatin by binding GGAA microsatellites whether by disturbing the TADs (Topological Associated Domains), which are epigenetics structures involved in the regulation of genes expression or by modulating chromatin loops, and thus, altering genes transcription (Showpnil *et al*, 2022). To go further, we were able to prove that among the 8 *KCNN1* transcripts described in the literature, one *KCNN1* transcript is drastically regulated by EWS-FLI1, the C transcript (NM_001386977.1), which is not the canonical *KCNN1* transcript, the A transcript (NM_002248.5). However, according to predictions of SK1 channel structures of the different variants, no radical change is observed in the structure of SK1 channel encoded by *KCNN1* C transcript, compared to the channel encoded by the *KCNN1* A transcript. Furthermore, *KCNN1* expression is not only regulated by EWS-FLI1 but also by EWS-ERG, confirming the ability of this rarer fusion protein to induce genes expressed only in ES tissues, as does EWS-FLI1 (Vibert *et al*, 2022). On the other hand, the overexpression of EWS-FLI1 in MSCs proves that the transcription of *KCNN1* C transcript requires the expression of the fusion protein, FLI1 not being sufficient to induce *KCNN1* transcription. The characterization of the SK1 channel lacking 3 amino acids in the calmodulin binding domain remain to be done in another study.

Regarding the role of *KCNN1* in a key function of tumor development, our results show that silencing of *KCNN1* expression decreases the ability of ES cells to form colonies and slows down the cell cycle. Historically, the involvement of potassium channels in cell proliferation was suggested by the work of DeCoursey *et al*., (DeCoursey *et al*, 1984), who demonstrated that voltage-gated potassium channels are the predominant channels in proliferative human T cells. The mechanisms by which potassium channels regulate cell proliferation are the subject of two consensus theories that can be linked: 1) oscillation of the membrane potential that participates in the control of cell volume and thus allows cells to pass the “cell volume checkpoint” for successful cell cycle progression, and 2) potassium efflux through potassium channel can lead to membrane hyperpolarization, which increases the driving force for calcium entry through a calcium-permeable channel at the plasma membrane. The increase in intracellular calcium concentration in turn promotes proliferation through calcium signaling (Huang & Jan, 2014). Although Fuest and colleagues did not recorded SK1 currents in ES cells (Fuest *et al*, 2022), our work shows that *KCNN1* silencing leads to membrane depolarization in a 2D and 3D cell models, and to a decrease of both SOCE and CCE, suggesting a mechanism of action according to the second hypothesis. In addition, the effect of SK1 channel activator on both membrane potential and calcium entries strongly suggests that *KCNN1* may express SK1 channel, which control plasma membrane potential of ES cells and thus calcium entries through calcium channels. The Ca^2+^ channel associated with this process channel remains to be identified in ES cells. In this context we can note that a study identified oncocomplexes of SK3 with calcium channels in various cancer cells, that control calcium entry and thus cancer migration and metastasis development (Potier-Cartereau *et al*, 2022b).

Regarding the expression of other SKCa channel genes in ES cells, our work shows that *KCNN3* expression increases after EWS-FLI1 silencing. These results suggest that cells that strongly express EWS-FLI1 would strongly express *KCNN1,* which in turn would stimulate cell proliferation. Conversely, cells with low expression of EWS-FLI1 would express *KCNN3* more strongly, which would participate in the stimulation of cell migration as observed in bone metastases (Potier *et al*, 2006). This hypothesis is consistent with the work of Franzetti *and al*., who showed that ES cells highly expressing EWS-FLI1 are characterized by highly active cell proliferation, and low EWS-FLI1 states where cells have a high propensity to migrate, invade and metastasize (Franzetti *et al*, 2017). In addition, Riggi *et al*. showed that the number of GGAA repetitions in microsatellites influences the regulation of EWS-FLI1-target genes (Riggi *et al*, 2014). Indeed, they proved that 4 repetitions – or more (within the limit of 20) – of GGAA activates EWS-FLI1-regulated genes, while genes repressed by EWS-FLI1 are associated with canonical ETS binding site (a single GGAA repetition) (Riggi *et al*, 2014). By a sequence analysis, only 2 GGAA repetitions are identified at the EWS-FLI1 binding site on *KCNN3* promoter, suggesting a repression of this gene by the fusion protein.

In conclusion, our results demonstrate the exclusive expression of *KCNN1* in ES patients identifying *KCNN1* as a novel biomarker of these tumors. Furthermore, our work dissects the regulation of transcription of the different *KCNN1* isoforms not only by EWS-FLI1 but also by EWS-ERG. Our work identifies the major role of *KCNN1* in the regulation of a key function in tumor development, cell proliferation, and suggests the role of calcium influx in these processes. Thus, our work identifies *KCNN1* as a potential therapeutic target in ES.

## Materials and Methods

### Cell culture and reagents

A673 (CRL-1598), SKES1 (HTB-86), RDES (HTB-166), TC71 (CVCL-2213), MHHES1 (CVCL-1411), EW24 (CVCL-1215) and EW3 (CVCL-1216) cells were a kind gift from Olivier Delattre (Curie Institute, France). ASP14 cell line, stably transduced with a doxycycline-inducible shRNA targeting EWS-FLI1, were also given by Olivier Delattre (Tirode *et al*, 2007). KHOS (CRL-1544), HOS-MNNG (CRL-1543), G292 (CRL-1423), MG63 (CRL-1427), SaOS2 (HTB-85) and U2OS (HTB-96) cell lines were purchased from ATCC (LGC Standards, Molsheim, France). ASP14, A673, SKES1, RDES, MHHES1, KHOS, HOS-MNNG, G292, MG63and U2OS cells were cultured in Dulbecco’s Modified Eagle’s Medium (DMEM, Lonza, Basel, Switzerland), supplemented with 10% Fetal Bovin Serum (FBS, Lonza); SaOS2 cells were cultured in DMEM supplemented with 15% FBS, whereas TC71, EW24 and EW3 cells were cultured in Rosewell Park Memorial Institute Medium (RPMI, Lonza) supplemented with 10% Fetal Bovin Serum (FBS, Lonza).

GW542573X was purchased from Tocris (Bristol, United Kingdom) and dissolved in Dimethyl Sulfoxide (DMSO, Sigma-Aldrich, St. Quentin Fallavier, France). Doxycycline was purchased from Sigma-Aldrich.

### Bioinformatics Analyses

R2: Genomics Analysis and Visualization Platform website provides free access datasets from patients biopsies. Multiple datasets were used to compare *KCNN1* expression (See Table EV1): Ewing sarcoma (“Tumor Ewing Sarcoma – Delattre – 117), Mesenchymal Stem Cells (MSCs, “Bonemarrow Mesenchymal stem cells” – Yamaguchi – 30), Glioma (“Tumor Glioma” – French – 284), Lungs (“Tumor Lung” – Chuang – 120), Pancreatic (“Tumor Pancreatic – Wu – 32), B-cell Lymphoma (“Tumor B-cell Lymphoma” – BAGS-subtypes – Boedker – 122), Melanoma (“Tumor Melanoma (Metastatic) – Matta – 87), Glioblastoma (“Tumor Glioblastoma – Hegi – 84), Medulloblastoma (“Tumor Medulloblastoma – Pfister – 223), Neuroblastoma (“Tumor Neuroblastoma – Hiyama – 51), Ovarian (“Tumor Ovarian” – Bowtell – 285), Breast (“Tumor Breast” – EXPO – 72), Colon (“Tumor Colon” – EXPO – 315), Kidney (“Tumor Kidney” – EXPO – 261), Prostate (“Tumor Prostate – EXPO – 72),

Osteosarcoma (“Tumor Osteosarcoma” – Kobayashi – 27), Rhabdomyosarcoma (“Tumor Rhabdomyosarcoma” – Barr – 58), Hepatocellular carcinoma (“Tumor Hepatocellular Carcinoma” – Llovet – 91), and Thyroid (“Tumor Thyroid – EXPO – 34). All of these datasets used the same u133p2 GeneChip and the MAS5.0 normalization.

Another dataset was used on the Kaplan Meier analysis, to compare the overall survival rate of Ewing sarcoma patients according to *KCNN1* expression: “Tumor Ewing Sarcoma (Core Exon) – Dirksen – 85. Differences between groups were calculated using the logrank test.

Databases GTEx (Genotype-Tissue Expression) and CCLE (Cancer Cell Line Encyclopedia) were respectively used to compare *KCNN1* expression between healthy tissues (GTEx version 8) and tumorous cell lines (CCLE - DepMap 22Q4).

### RNA-Seq Analyses

RNA-seq was performed for A673, EW3, EW24, MHHES1, RDES, SKES1, and TC71 ES cell lines, on KHOS, HOS-MNNG, G292, MG63, U2OS and SaOS2 OS cell lines and on A673 shRNA 908 treated or not with doxycycline during 7 days (GSE229906).

#### Gene-level quantification

RNA-Seq data aligned on the genome, libraries were prepared at Active Motif Inc. using the Illumina TruSeq Stranded mRNA Sample Preparation kit, and sequencing was done on the Illumina NextSeq 500 as 42-nt long-paired end reads. Fastp (v. 0.19.5 (Chen *et al*, 2018)) was used to filter low quality reads (Q > 30) and trim remaining PCR primers. Read mapping against the human genome (hg19) was done using HISAT2 (v 2.1.0 (Kim *et al*, 2019)) and fragment quantification was done using stringtie (v 2.1.1(Pertea *et al*, 2015)). Differential gene expression analysis was performed using DESeq2 package (Love *et al*, 2014). A gene is considered significantly differentially expressed if its log2 fold change was 1 or less than -1, and the false discovery rate (FDR) was less than 0.05.

#### Transcript-level quantification

Here we also included RNA-Seq data for ASP14 from the Gene Expression Omnibus (GEO with the accession: GSE164373) without induction (day 0) and with induction of shRNA targeting *EWS-FLI1* (day 7) performed in triplicates. The reads were filtered with fastp (v. 0.20.1) (-q 30 -u 40). They were then aligned to the transcriptome associated with RefSeq (GRCh38.p14) via kallisto (v. 0.46.2) (Bray *et al*, 2016). Counts were normalized with DESeq2 (v. 1.30.1) (Love *et al*, 2014). Transcripts were considered differentially expressed if their expression levels between conditions were affected by at least 2-fold, and had an adjusted p-value less than 0.05 (|log2 FC| > 1 & adj.p.value < 0.05), according to a Wald test.

### ChIP-Seq Analyses

ChIPseq data were acquired from GEO with accessions GSE176400, GSE129155 and GSE94278. GSE176400 contains ChIPseq from 18 Ewing sarcoma cell lines targeting FLI1 or ERG depending on the fusion protein present in the cell line; histone marks H3K4me3, H3K27me3, and H3K27ac; and an input serving as a control. GSE129155 contains ChIPseq against the H3K27ac mark in the ASP14 cell line without (day 0) or with (day 7) EWS-FLI1 knock-down, and ChIPseq against FLI1 in the same cell line starting at the end of the shRNA induction (day 7), and then on days 9, 10, 11, 14, and 17, in order to observe the effects of gradual reappearance of EWS-FLI1. These data are supplemented with an input from day 0 that served as a control. GSE94278 contains Bigwig files of normalized ChIP-Seq values of MSCs transfected with a control plasmid, FLI1 plasmid, and EWS-FLI1 plasmid. A few corrupted reads (nucleotide sequence length different from the phred score) were removed using Awk (v. 4.0.2) (Aho *et al*, 1979). ChIPseq data were all filtered with fastp (v. 0.23.2) (Chen *et al*, 2018) by removing reads with more than 40% of bases with a phred score below 30 and removing PCR adapters if detected (-q 30 -u 40). Reads were aligned with BWA-MEM (v. 0.7.17) (Li & Durbin, 2009) against the GRCh38.p14 reference genome “analysis set” version from NCBI. After conversion to BAM format and sorting of reads according to their coordinates with samtools (v. 1.16.1)(Danecek *et al*, 2021), peaks were detected with Macs (v. 2.2.7.1) (Zhang *et al*, 2008) using the controls associated with each sample. Sequencing depths of the alignment files to construct the plots were retrieved through python scripts using Pysam (v. 0.20.0) (https://github.com/pysam-developers/pysam). Macs2 sequencing depths and peaks coordinates plots were constructed with ggplot2 3.4.0 (Wickham, 2009), and transcript representations with gggenes 0.4.1 (https://github.com/wilkox/gggenes) from the RefSeq GTF file associated with the reference genome used. These graphs were assembled with Cowplot 1.1.1 (https://github.com/wilkelab/cowplot). As for number of microsatellite repeats at the EWS-FLI1 or EWS-ERG binding site, the consensus sequence of the region was obtained with bcftools mpileup 1.16 (Danecek *et al*, 2021) by keeping only mutations present at least 5 times, then the counting was done by TandemRepeatFinder 4.09.1(Benson, 1999) by strongly penalizing the mismatches and indels.

### Protein structure predictions

Protein structure predictions were obtained using the AlphaFold2_mmseqs2 notebook in ColabFold 1.5.2 (Mirdita *et al*, 2022), and were then displayed using Mol* (Sehnal *et al*, 2021).

### RT-qPCR (Reverse Transcription – quantitative Polymerase Chain Reaction)

RNA from cell lines were extracted using the NucleoSpin RNA Plus Kit (Macherey-Nagel, Duren, Germany) and dosed with the Nanodrop 1000 spectrophotometer (ThermoFischer, Illkirch, France). Then, the Reverse Transcription (RT) was performed from 1 µg of RNA with the “Maxima H Minus First Strand cDNA Synthesis Kit, with dsDNase” kit (ThermoFischer) according to the supplier’s protocol. The quantitative Polymerase Chain Reaction (qPCR) was realized with 20 ng of cDNA using SYBR Select Master Mix (ThermoFischer), and primers listed on the Table I. Real-time PCR was performed on a CFX 96 real-time PCR instrument (Biorad, Richmond, CA, USA). Target gene expression was normalized to HPRT, B2M or ActinB levels in respective samples and the Cycle threshold (Ct) method was used to measure the relative quantification of target mRNAs.

### Western blot

Cells were lysed in Sodium Dodecyl Sulfate (SDS) buffer (SDS 1%; pH 7,4; Tris 10 mM Na_3_VO_4_ 1 mM) supplemented with 1:500 diluted benzonase (Sigma-Aldrich). Protein concentration was determined using the Bicinchoninic acid assay (BCA, Sigma-Aldrich). Proteins were then denaturated with Laemmli Buffer (Tris pH 6.8 (200mM), 8% SDS, 40% de Glycerol, 0.04% bromophenol blue, 20% β-mercaptoethanol) at 95°C for 5 minutes, before being loaded in SDS Polyacrylamide Gel. After electrophoresis separation, proteins were transferred to PolyVinyliDene Fluoride 0.45 µm membranes (Immobilon-Fl, Merck Millipore, Darmstadt, Germany). Membranes were saturated 1 hour with a Tris-Buffered Saline blocking solution (TBS blocking solution, Odyssey Blocking Buffer, Li-Cor, Nebraska, USA) at room temperature and were incubated overnight at 4°C with primary antibodies diluted in previous blocking buffer. The antibodies used for western blotting were Aurora A (1:1000, CST #91590S (Li *et al*, 2022)), p21 (1:1000, CST #2947S (Schloesser *et al*, 2023)) and β-tubulin (1:2000, CST #186298S (Mabrouk *et al*, 2022)). After wash, membranes were incubated with a secondary fluorescent antibody (1:20000, Donkey anti-rabbit IRDye 680RD #926-68073 ; 1:20000 Goat anti-mouse IRDye 800CW #926-32210) and diluted in the previous blocking buffer for 1 hour in the dark at room temperature. The labelled proteins were detected by fluorescent detection, using the Odyssey FC Device (Cambridge, UK) and the Image Studio software.

### siRNA transitory transfection

Twenty-four hours after seeding, A673, MHHES1, RDES and SKES1 cells at 50-60% confluency were transfected with a pool of 5 siRNAs targeting *EWS-FLI1* (siRNA EWSR1, ON-TARGET, SmartPool, 50 µM, Dharmacon, Horizon, Perkin-Elmer), targeting *KCNN1* (SK1 siRNA (h), sc-36494, 10 µM, SantaCruz, Dallas, TX, USA) or a pool of 5 control siRNAs coming from Dharmacon (ON-TARGET plus, Non-Targeting Pool 50 µM, Dharmacon, Horizon, Perkin-Elmer) or SantaCruz (Control siRNA-A, sc-37007, 10µM, SantaCruz) at a final concentration of 0,03 µM, using Opti-MEM medium and the Lipofectamine RNAiMax (ThermoFischer) according to the supplier’s recommendations. Sixteen hours after transfection, cells are-re-seeded for proliferation assays, and fourty-eight hours after transfection, RNAs and proteins are harvested for RT-qPCR and Western blot respectively.

### shRNA stable transduction

Two different cell lines, A673 and SKES1 were transduced with 3 doxycycline-inducible shRNAs targeting *KCNN1* (V3IHSMCG_7156511; V3IHSMCG_4737908 and V3IHSMCG_9443576; Dharmacon) or a doxycycline-inducible shRNA non-targeting control (Non-targeting Control mCMV – Turbo GFP, VSC6584, Dharmacon) at a Multiplicity of Infection 1 (MOI 1). After the transduction, cells were selected with puromycin (1,25 µg/mL, Sigma-Aldrich) for seven days. The expression of *KCNN1* was measured by RT-qPCR after 72 hours or 7 days of doxycycline treatment (0,5 µg/mL). Before any experiment, cells were pre-treated with doxycycline (Sigma-Aldrich) for 72 hours or 7 days at 0,5 µg/mL and the treatment was maintain afterwards.

### Membrane depolarization measures

Concerning DiBAC4(3) measures, cells were plated in 96 well plates at 20000 cells per well 24 hours before the experiment. Extemporaneously, 0,18% glucose was added to Physiological Salin Solution (PSS, 137 mM NaCl, 5,6 mM KCl, 1 mM MgCl_2_, 2 mM CaCl_2_, 0,42 mM Na_2_HPO_4_ (anhydrous), 0,44 mM Na_2_HPO_4_ (hydrated), 10 mM 4-(2-hydroxyethyl)-1-piperazineethanesulfonic acid (HEPES), 4,17 mM NaHCO_3_, pH 7,4. DiBAC4(3) stock solution was prepared following Thermofischer recommendations. Adherent cells were loaded for 15 minutes à 37°C with the 1 µM DiBAC4(3) PSS. Fluorescence was measured at 520 nm using the FlexStation Microplate Reader (Molecular Devices, San Jose, CA, USA), in response to an excitation wavelength of 485 nm. Then, a second fluorescence measure was taken, after a 10 minute-incubation of the cells with 100 µL of DiBAC4(3) 1µM.

For membrane depolarization measures on spheroids, they were firstly formed through use of the scaffold-free method cell suspensions (100 µL of cell suspensions in DMEM supplemented with 10% FBS were added at a density of 5.10^4^/mL). Cell suspensions were added to Ultra-Low Attachment microplates (Corning, Glendale, AZ, USA) and the plates were incubated stationary in standard cell culture conditions for 72 hours. To measure spheroids membrane potential, as described previously (Le Guennec *et al*, 2002), glass microelectrodes were backfilled with 3 M KCl and then connected to an impedance adapter (HS 170, Biologic, Seyssinet-Pariset, France), linked to an amplifier (VF-180, Biologic). The resistances of the microelectrodes range between 15 and 20 MΩ.

### Clonogenicity assays

Five hundred A673 or SKES1 cells transduced with shRNAs targeting *KCNN1* or shRNA non-targeting control were seeded in 6-well plates after 72 hours or 7 days of doxycycline treatment (0,5 µg/mL, Sigma). Cells were then incubated for 8 to 10 days under standard conditions (humidified incubator at 37°C and 5% CO2). Once colonies were detectable under a microscope, cells were fixed with 1% glutaraldehyde and stained by crystal violet. The number of colonies was then counted with the Cell Counter from Fiji software and their surface was measured with a macro created by the Cellular and Tissular Imaging Core Facility of Nantes University (MicroPICell, Nantes, France).

### Viability assays

After siRNA transfection, viability assays were performed on re-seeded cells, using the Omni device from CytoSMART (Axion Biosystems, Atlanta, GA, USA). Cell viability was monitored for 72 hours in time-lapse, with a scan every 4 hours. Data were collected on the CytoSMART Cloud and the analyses were performed on the same software.

### Thymidine synchronization

After a 72 hours pre-treatment with doxycycline (0,5 µg/mL), A673 cells transduced with shRNAs targeting *KCNN1* or shRNA non-targeting control were seeded in ø 60mm Petri boxes. Once the cells were adhered, they were treated with 2 mM thymidine (Sigma-Aldrich) for 18 hours, washed twice with PBS and left in DMEM 10% FBS, supplemented or not with doxycycline, for 8 hours. Then the cells were treated a second time with 2 mM thymidine for 16 hours and washed twice with PBS before being left in DMEM 10% FBS supplemented or not with doxycycline for 12 hours. Cell distribution was observed by flow cytometry in Platform Cytocell in Nantes University (FACSymphony A5, BD, Franklin Lakes, USA), after 0 hour, 6 hours, 8 hours, 10 hours and 12 hours or not synchronized. In addition, proteins were extracted for western blot analyses.

### Cell Cycle Analysis

After thymidine synchronization, cells were washed with PBS, trypsinized and fixed with 70% ethanol. Cells were washed again with PBS, centrifuged and resuspended in a phospho-citrate buffer (Na_2_HPO_4_ 0.2 M, citric acid C_6_H_8_O_7_ 0.1 M, pH 7.5) for 30 minutes at room temperature. Cells were centrifuged and resuspended in PBS, 0.12% Triton, 0.12 mM EDTA and 100 µg/mL RNAse A (Promega A797c, Promega) and incubated 30 minutes at 37°C. Then Propidium Iodide (50 µg/mL) was added for 20 minutes in the dark at 4°C. Cell distribution was observed by flow cytometry in Platform Cytocell in Nantes University (FACSymphony A5). Data were analyzed with Multicycle software.

### Store operated Ca^2+^ entry measurement by Fura-2 AM

As already published (Ibrahim *et al*, 2021), cells were plated in 96 well plates at 20000 cells per well 24 h before the experiment. Adherent cells were loaded for 45 min at 37°C with the ratiometric dye Fura2-AM (5 μM) then washed by PBS solution supplemented with Ca^2+^.

During the experiment, cells were incubated with Physiologic Saline Solution PSS Ca^2+^ free solution and treated by 2 μM thapsigargin (TG, T7458, Life Technologies, ThermoFisher) to deplete intracellular store of Ca^2+^. Calcium entry was stimulated by injection of 2 mM of CaCl2. Fluorescence emission was measured at 510 nm using the FlexStation-3 (Molecular Devices, San Jose, CA, USA) with excitation at 340 and 380 nm. Maximum of fluorescence (peak of Ca^2+^ influx F340/F380) is measured and compared to normal condition.

### Constitutive Ca^2+^ entry measurement by Mn^2+^ quenching

As already published (Ibrahim *et al*, 2021), cells were plated in 96 well plates at 20000 cells per well 24 h before the experiment. Adherent cells were loaded for 45 min at 37°C with the ratiometric dye Fura2-AM (5 μM) then washed by PBS solution supplemented with Ca^2+^. During the experiment, cells were incubated with PSS Ca^2+^ free solution and treated by 0.9 mM of Manganese Mn^2+^ (manganese chloride, Sigma-Aldrich). Fluorescence emission was measured at 510 nm using the FlexStation-3 with excitation at 360 nm.

### Statistical Analyses

Statistical analyses were performed using GraphPad Prism Software. Comparison of SKCa (*KCNN1*, *KCNN2* and *KCNN3*) gene expression in patients suffering from Ewing sarcoma was performed by a Brown-Forsythe and Welch Anova and a Games-Howell multiple comparisons tests. The comparison of *KCNN1* gene expression between ES patients and MSCs coming from healthy donors was performed by an unpaired t-test and a Welch correction. The comparison of *KCNN1* gene expression between multiple tumors was performed by a Kruskal-Wallis test and a Dunn’s multiple comparisons test. For *in vitro* experiments, to compare two groups, Mann-Whitney tests were used and Anova tests were performed when N > 30 and normality test was passed. Wilcoxon tests were used during cell cycle assays. α=0.05 and p<0.05 is considered statistically significant: * p<0.05 ; ** p<0.01 ; *** p<0.001 ; **** p<0.0001.

## Data availability Section

The datasets and computer code produced in this study are available in the following databases:

- RNA-Seq data: Gene Expression Omnibus GSE 229906
- Modeling computer scripts: GitHub (https://github.com/EpistressLab/KCNN1_ewing_sarcoma)

## Acknowledgements

This work was supported by INCA (# 2018-151), Ligue contre le cancer (CD 41, 44, 49, 56 et 85), M la vie avec Lisa, Imagine for Margo, Societé Française de lutte contre les cancers et les leucémies de l’enfant et de l’adolescent (SFCE), l’étoile de Martin, Enfants Cancers Santé. We thank the Cytometry Facility « Cytocell » from Nantes for their expert technical assistance » (Cytocell).

## Author contributions

**Maryne Dupuy**: Conceptualization, Data curation, Formal analysis, Investigation, Methodology, Validation, Writing original draft. **Maxime Gueguinou**: Conceptualization, Data curation, Formal analysis, Investigation, Writing review and editing. **Anaïs Postec**: Data curation, Formal analysis, Investigation, Software. **Régis Brion**: Investigation. **Robel Tesfaye**: Formal analysis, Writing review and editing. **Mathilde Mullard**: Writing review and editing. **Laura Regnier**: Formal analysis. **Jérôme Amiaud**: Investigation. **Marie Potier-Cartereau**: Conceptualization, Validation. **Aurélie Chantôme**: Conceptualization, Validation. **Bénédicte Brounais-Le Royer**: Writing review and editing. **Marc Baud’huin**: Writing review and editing. **Steven Georges**: Writing review and editing. **François Lamoureux**: Writing review and editing. **Benjamin Ory**: Writing review and editing. **Olivier Delattre**: Conceptualization, Writing review and editing. **Françoise Rédini**: Conceptualization, Writing review and editing. **Christophe Vandier**: Conceptualization, Formal analysis, Writing review and editing. **Franck Verrecchia**: Conceptualization, Formal analysis, Fundig acquisition, Project administration, Ressources, Supervision, Validation, Writing original draft.

## Disclosure and competing interests statement

The authors declare that they have no competing interests.

## The Paper Explained

### Problem

Ewing sarcoma (ES) is the second most common primary malignant bone tumor in children and adolescents. Patient survival has not increased during the past decade, and remaining too low for high-risk patients, with a 5-year survival rate of 30%. This tumor is characterized by a chromosomic translocation, which leads in 85% of cases to the formation of the EWS-FLI1 fusion protein and in 10% of cases to the formation of the EWS-ERG fusion protein, both acting as an aberrant transcription factor for many oncogenes. More and more studies have emerged on the implication of ion channels in tumorigenesis. Indeed, potassium channels have been found to be involved in the regulation of cell proliferation, migration or apoptosis in many cancers. More precisely, small conductance calcium-activated potassium channels SK2 and SK3 were respectively shown to be involved in melanoma and in the formation of bone metastasis coming from breast cancer. In ES, SK1 has been observed as overexpressed in ES cell lines, but its expression in patients biopsies and its involvement in ES tumorigenesis has not been established.

## Results

Based on data from tumor-suffering patients, we observed exclusive overexpression of *KCNN1*, the gene encoding the SK1 channel, in ES patients, and except for brain tissues, no other healthy tissues express *KCNN1*. At a cellular level, we also demonstrated regulation of *KCNN1* by the fusion proteins EWS-FLI1 and EWS-ERG, which bind to specific regions of the *KCNN1* promoter, called GGAA microsatellites. *KCNN1* silencing in ES cell lines resulted in an impairment in cell proliferation, with a slowdown of the cell cycle. Our results also showed that *KCNN1* expression modulates membrane potential and calcium flux in ES cells. Based on our data, we propose that SK1 is directly regulated by EWS-FLI1 or EWS-ERG and that it is involved in a key function of ES tumor development, the ES cell proliferation.

### Impact

Our work showed for the first time the restricted overexpression of *KCNN1* in ES patients. We also proved the involvement of SK1 channel in a key process of ES tumorigenesis, the cell proliferation via the regulation of cells progression in the cell cycle. These data could allow SK1 to be considered as a potential therapeutic target in ES.

## For more information

INCA (https://www.e-cancer.fr)

Ligue contre le cancer (https://www.liguecancer44.fr) M la vie avec Lisa (http://www.mlavieaveclisa.com) SFCE (https://sf-cancers-enfant.com)

Imagine for Margo (https://imagineformargo.org)

Enfant Cancer Sante (https://www.enfants-cancers-sante.fr)

## Figure legends

**Table 1:**
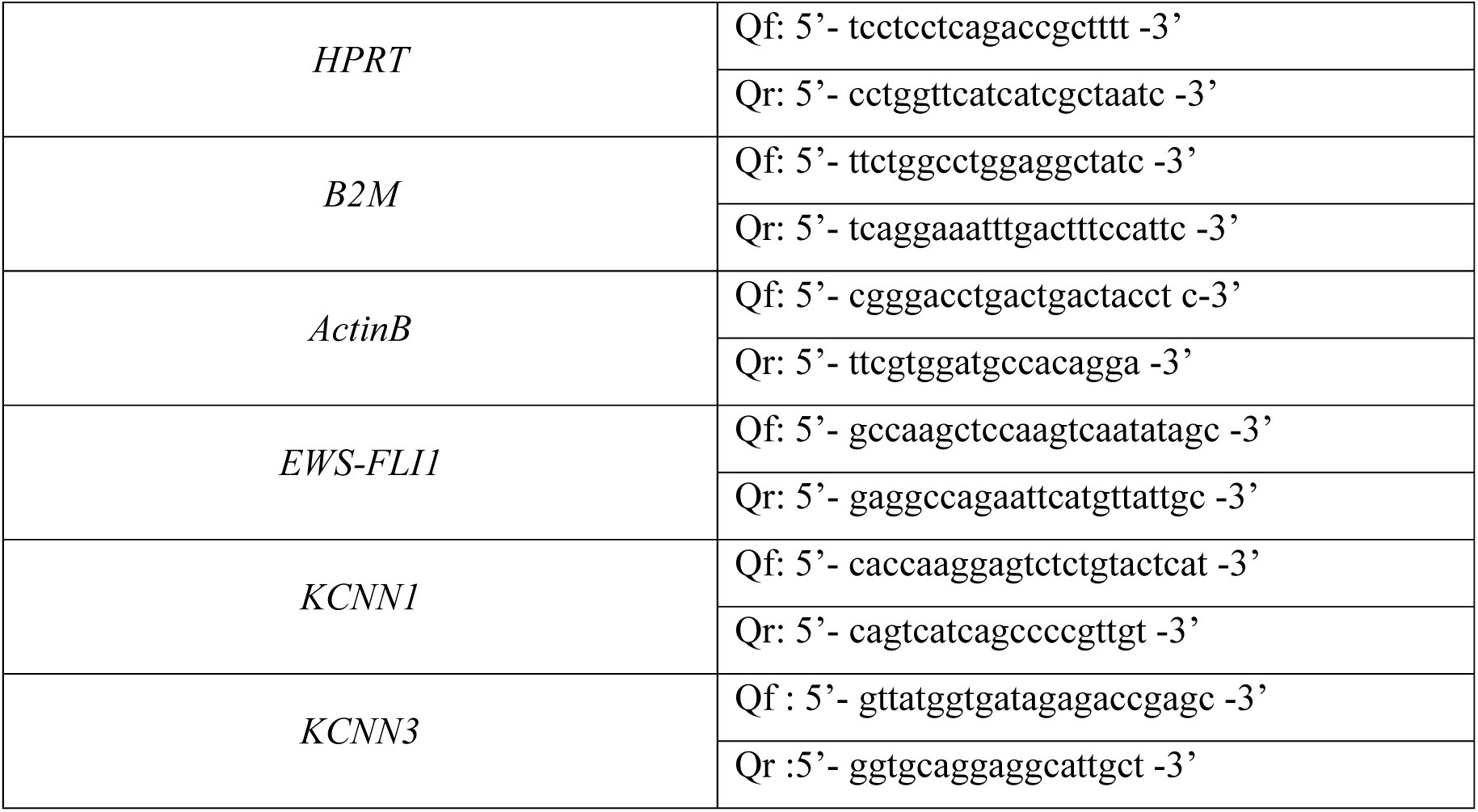
Primers used for qPCR experiments

## Expanded View Figure legends

**Figure EV1: *KCNN2* and *KCNN3* are ubiquitously expressed in patients’ biopsies.**

A Gene expression (Log2 normalized expression) of *KCNN2* in patients suffering from different tumors.

B Gene expression (Log2 normalized expression) of *KCNN3* in patients suffering from different tumors.

Data information: The boxes indicate the first and third quartiles - and the median inside – of genes expression in Log2 Fold change. The whiskers indicate the fifth and the ninety-fifth percentiles.

**Figure EV2: Multiple *KCNN1* transcripts are expressed in ES cell lines.**

A Normalized expression of *KCNN1* transcripts (FPKM) in EW3, EW24, MHHES1, RDES, SKES1 and TC71 cell lines (N=3).

B Predictions of structures of SK1 channel encoded by KCNN1 A, B or C transcripts. Pink structures on transcripts A and B correspond to the changes compared to the transcript C.

**Figure EV3: EWS-FLI1 binds at GGAA microsatellites near *KCNN1* promoters in multiple ES cell lines and regulate *KCNN3* expression in ASP14 cell line.**

A H3K4me3, H3K27ac and FLI1 ChIP-seq data with quantitative value of peaks around *KCNN1* promoters and sequences of EWS-FLI1 binding sites in 13 ES cell lines.

B Normalized and relative *KCNN3* gene expressions in ASP14 cell line after doxycycline treatment (1 µg/mL). Bars indicate mean ± SD of relative and normalized *KCNN3* mRNA expression determined by RT-qPCR (N=5).

C FLI1 ChIP-Seq of ASP14 cell line with quantitative value of FLI1 peaks around *KCNN1* promoters’ areas, at day 7 of doxycycline treatment, and 3 days (day 10), 4 days (day 11), 7 days (day 14) and 10 days (day 17) after discontinuation of doxycycline treatment.

**Figure EV4: *KCNN1* C transcript is regulated in shRNA 908 cells and its expression regulates cell proliferation in MHHES1 cell line.**

A Normalized expression of *KCNN1* transcripts (FPKM) in A673 cell line transduced with shRNA 908 after 72 hours of doxycycline treatment (0,5 µg/mL) (N=2).

B Verification of *KCNN1* extinction in A673 cell line stably transduced with doxycycline-inducible shRNA control cells (shRNA CT) or 3 shRNA cells targeting *KCNN1* (shRNA 511, shRNA 908 and shRNA 576) after 7 days of treatment with doxycycline (0,5 µg/mL). Bars indicate mean ± SD of relative and normalized *KCNN1* mRNA expression obtained after RT-qPCR (N=3).

C Viability assay during 64 hours on MHHES1 after *KCNN1* silencing (N=3).

D p21 and Aurora A expression kinetics after the thymidine double block, normalized to β-tubulin.

Data information: *: p < 0,05 (Mann-Whitney tests).

**Figure EV5: *KCNN1* expression modulates calcium flux in SKES1 cell line.**

A Effect of *KCNN1* knockdown on SOCE in SKES1 cell line stably transduced with shRNA control (shRNA CT) or a shRNA targeting *KCNN1* (shRNA 908). Left panel: Fluorescence measurement of SOCE induced by thapsigargin (TG). Right panel: Fluorescence measurement of SOCE induced by TG in shRNA *KCNN1* conditions compared to control on the relative fluorescence of shRNA CT cells loaded with Fura2-AM (N=26 for shRNA CT and N=29 for shRNA 908).

B Effect of *KCNN1* knockdown on constitutive calcium entries using Mn2+ quenching assay. Left panel: Effect of *KCNN1* knockdown on Mn2+ quenching rate. Right panel: Effect of *KCNN1* knockdown on normalized Mn2+ quenching slope (N=21 for shRNA CT and N=23 for shRNA 908).

Data information: **: p < 0,01 ; ***: p < 0,001 (Mann-Whitney tests).

---

**Table EV1:** Cohorts used on R2 platform.

| Sample | Author | Platform | Normalization | Number of samples | Accession | Release date |
| --- | --- | --- | --- | --- | --- | --- |
| Ewing sarcoma | Delattre | u133p2 | MAS5.0 | 117 | GSE34620 | 01.01.2000 |
| MSCs | Yamaguchi | u133p2 | MAS5.0 | 30 | GSE7637 | 06.06.2008 |
| Glioma | French | u133p2 | MAS5.0 | 284 | GSE16011 | 01.01.2000 |
| Lung | Chuang | u133p2 | MAS5.0 | 120 | GSE19804 | 30.01.2011 |
| Pancreatic | Wu | u133p2 | MAS5.0 | 32 | GSE32676 | 08.10.2011 |
| B-cell<br>Lymphoma | Boedker | u133p2 | MAS5.0 | 122 | GSE56315 | 24.03.2015 |
| Melanoma | Matta | u133p2 | MAS5.0 | 87 | GSE7553 | 28.05.2008 |
| Glioblastoma | Hegi | u133p2 | MAS5.0 | 84 | GSE7696 | 01.01.2000 |
| Medulloblastoma | Pfister | u133p2 | MAS5.0 | 223 |  | 27.03.2017 |
| Neuroblastoma | Hiyama | u133p2 | MAS5.0 | 51 | GSE16237 | 01.09.2009 |
| Ovarian | Bowtell | u133p2 | MAS5.0 | 285 | GSE9891 | 01.03.2008 |
| Breast | EXPO | u133p2 | MAS5.0 | 72 | GSE2109 | 15.01.2005 |
| Colon | EXPO | u133p2 | MAS5.0 | 315 | GSE2109 | 15.01.2005 |
| Kidney | EXPO | u133p2 | MAS5.0 | 261 | GSE2109 | 15.01.2005 |
| Prostate | EXPO | u133p2 | MAS5.0 | 72 | GSE2109 | 15.01.2005 |
| Osteosarcoma | Kobayashi | u133p2 | MAS5.0 | 27 | GSE14827 | 01.02.2010 |
| Rhabdomyo-<br>sarcoma | Barr | u133p2 | MAS5.0 | 58 | GSE66533 | 01.06.2016 |
| Hepatocellular<br>carcinoma | Llovet | u133p2 | MAS5.0 | 91 | GSE9843 | 22.06.2008 |
| Thyroid | EXPO | u133p2 | MAS5.0 | 34 | GSE2109 | 15.01.2005 |

## Notes

### Competing Interest Statement

The authors have declared no competing interest.

